# Label-free horizontal EMSA for analysis of protein-RNA interactions

**DOI:** 10.1101/825679

**Authors:** William Perea, Nancy L. Greenbaum

## Abstract

We describe a method to analyze the affinity and specificity of interactions between proteins and RNA using horizontal PAGE under non-denaturing conditions. The method permits tracking of migration of anionic and cationic biomolecules and complexes toward anode and cathode, respectively, therefore enabling quantification of bound and free biomolecules of different charges and affinity of their intermolecular interactions. The gel is stained with a fluorescent intercalating dye (SYBR^®^Gold or ethidium bromide) for visualization of nucleic acids followed by Coomassie^®^ Brilliant Blue R-250 for visualizations of proteins; the dissociation constant is determined separately from the intensity of unshifted and shifted bands visualized by each dye. The method permits calculation of bound and unbound anionic nucleic acid and cationic protein components in the same gel, regardless of charge, under identical conditions, and avoids the need for radioisotope or fluorescent labeling of either component.

## Introduction

The electrophoretic mobility shift assay (EMSA) is a simple but powerful method for characterization of the interaction of protein and RNA. The assays use polyacrylamide gel electrophoresis (PAGE) under non-denaturing conditions and buffers at or near neutral pH. The migration properties for free and bound biomolecules in an electric field are quantified to determine affinity and relative specificity of the interaction [1]. Conventional methods require the pre-labeling of the nucleic acid with a radioisotope or a fluorescent dye, a process that can be tedious and expensive [2]. However, many RNA-binding proteins (RBPs) have a high isoelectric point (pI) [3] and are thus positively charged at pH values of most electrophoresis buffers, in which case migration toward the cathode during electrophoresis in a standard vertical gel results in loss of the protein or protein-nucleic acid complex into buffer or the need to electrophorese the components at an unphysiological pH.

In this report we present an EMSA method for analysis of RBPs and their complexes with RNA. The method makes use of non-denaturing PAGE in a horizontal format, in which the wells are placed in the middle of the gel. Samples in loading buffer with mixed dyes are loaded onto pre-equilibrated gels and electrophoresed at 4 °C and constant voltage. After electrophoresis the gels are orthogonally stained, first with a fluorescent intercalating dye, *e.g*. ethidium bromide (EtBr) or SYBR^®^Gold, depending on desired sensitivity. After visualization of the RNA bands, the gels are stained with Coomassie Brilliant Blue R-250. The advantage of this approach is the potential to measure relative amounts of bound and unbound anionic nucleic acid and cationic protein components in the same gel, regardless of charge, and under identical conditions, without the need for labeling of biomolecular samples. A manuscript reporting the use of an earlier version of this method was published previously [4].

## Materials and methods

### Reagents

Proteins were expressed and purified in our laboratory, with the exception of lysozyme and BSA, which were purchased from Sigma (St. Louis, MO, USA) and Thermo Scientific (Rockford, IL, USA), respectively. Prior to the analysis, purity and homogeneity was confirmed by SDS PAGE and non-denaturing PAGE. Short RNAs (up to 20 nucleotides) were purchased from Dharmacon Inc.; RNA oligomers >20 nucleotides were prepared by *in vitro* transcription in the laboratory and purified by denaturing PAGE and electroelution. Acrylamide and bis-acrylamide were purchased from Acros Organics (Fair Lawn, NJ, USA). Coomassie Brilliant Blue R-250 was purchased from Fisher Scientific (Fair Lawn, NJ, USA). SYBR™Gold was purchased from Invitrogen (Eugene, OR, USA). All other reagents were of analytical or molecular biology grade. Buffers were prepared with distilled, deionized water (ddH_2_O) and filter sterilized prior to use. Individual binding buffers conditions are described in each figure legend.

### Non-denaturing gel electrophoresis

Separating gels were 5% polyacrylamide (diluted from a 40% stock of acrylamide:bis-acrylamide, 38:2) prepared in a buffer composed of 30 mM MOPS/25 mM histidine, pH 6.5. To speed polymerization, the gel mix (50 mL) was heated in a microwave oven for 30 sec, after which 200 μL 1% APS was added, followed by 50 μL TEMED. The solution was then poured immediately into a UV-transparent gel tray caster (100 mm x 70 mm x 20 mm), and a multi-tooth comb (1mm thick, with eight teeth for these experiments), was placed across the center of the gel. After polymerization of gel (final thickness 10 mm), the gel tray was assembled in a submerged horizontal electrophoresis system (Bio-Rad Mini-Sub Cell^®^ System) and equilibrated with running buffer (same as above) at 4 °C at 100 V for at least one hour. Samples were prepared in binding buffer (20mM NaPi pH 6.5, 100mM NaCl, 1mM DTT) and mixed with 1/6 volume 6x loading buffer (30% glycerol, 0.25% xylene cyanol, 0.25% bromophenol blue) prior to loading, after which they were electrophoresed at 4 °C at 100 V for 60-90 min.

### Staining and imaging of RNA oligomers

After electrophoresis, the gel in the tray was stained with 100 mL 2X SYBR™ Gold (diluted from 10,000X concentrate, as purchased, in DMSO) and protected from light for one hour (the relatively long incubation because of the thickness of the gel). For larger RNA or where lesser sensitivity is acceptable, staining with EtBr (1 μg/mL) provides sufficient intensity. RNA bands were visualized by trans-illumination at 302 nm and photographed with a UVP GelDoc-It™ Imaging System equipped with a Gel HR Camera. Optimal photographic conditions were determined empirically.

### Staining and imaging of protein

After staining with the fluorescent dye, the same gel was stained to visualize protein following a rapid-staining protocol by Dong et al. [5]. Briefly, the gel was transferred carefully to a plastic tray filled with 150 mL ddH_2_O, and heated in a microwave oven for 60 sec to help remove the fluorescent dye. After the water was discarded, the gel was stained with 100 mL Coomassie Blue (0.25 mg/mL Brilliant Blue R-250 in ddH_2_O) and again heated in a microwave oven for 60 sec. The staining solution was discarded and the gel washed with 150 mL ddH_2_O and heated in a microwave oven for 60 sec; this procedure was repeated twice more. Protein migration was visualized on a UV/White converter plate and photographed with a Gel HR Camera as part of a UVP GelDoc-It™ Imaging System. For additional sensitivity to assist visualization of faint bands, we used the classic method [], by which the gel was fixed with 100 mL 10% acetic acid, 50% methanol for one hour, stained with 0.025% w/v Coomassie Brilliant Blue R-250 in 10% acetic acid overnight, and destained with 10% acetic acid for 2 h. Further distaining was obtained by soaking the gel in ddH_2_O overnight.

### Data Analysis

To determine the equilibrium dissociation constant (K_d_) of the biomolecular interaction, intensities of RNA and protein bands were quantified using the 1D Analysis plug-in module of VisionWorks LS software (UVP, Upland, CA, USA). The band intensities and volumes were quantified for each shifted and unshifted band and plotted versus the concentration expressed in micromolar (μM) units. The K_d_ values were obtained by non-linear regression (hyperbola), in this case using the model of one-site binding (see Results for rationale), from GraphPad Prism version 8.0 for windows (GraphPad Software, San Diego, California USA, www.graphpad.com):

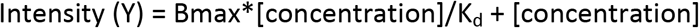

where Y represent the intensity or intensity volume of shifted band and Bmax the maximum intensity.

## Results

To illustrate the advantage of the horizontal gel configuration with the wells centered between the anode and cathode, Figure 1 shows the migration of two proteins, anionic BSA (pI = 4.8) and cationic lysozyme (pI = 11.3) at pH 6.5, migrating in opposite directions.

**Figure 1.**
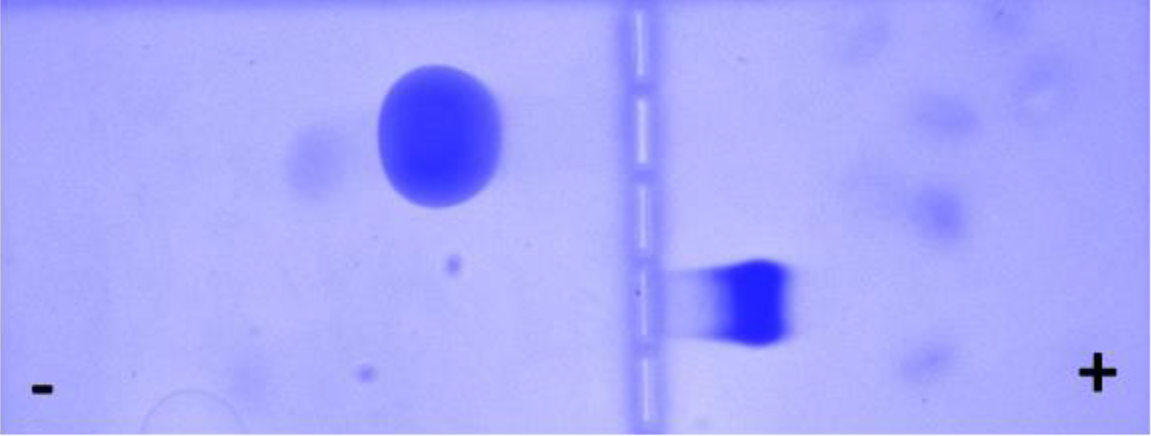
Horizontal nondenaturing PAGE displaying migration of anionic (BSA) and cationic (lysozyme) proteins. PAGE was performed by loading 15 μL of 2 μg/μL protein solution mixed with 3μL 6X mixed dyes solution on the wells in the middle of a gel (5% Acrylamide-N,N-bisacrylamide, 38:2). After running with non-denaturing buffer (30 mM MOPS-25 mM histidine pH 6.5), at constant voltage (100V) for 1 hour, at 4 °C, the gel was stained with Coomassie Blue R-250 according to the protocol published by Dong et al. [5]. The figure displays migration of BSA (pI 4.8) toward the anode and migration of lysozyme (pI 11.3) toward the cathode.

To demonstrate the utility of the method for analysis of protein-RNA interactions, we measured the affinity of the bimolecular interaction of a soluble protein containing a single RNA recognition motif (RRM), the human splicing factor SF3b14a/p14 (hereinafter p14), with a short RNA oligomer representing the spliceosome’s branch site region of an intron [4]. For each EMSA assay, we mixed 30 μM p14 with RNA (concentrations ranging from 9-90 μM). After electrophoresing on a horizontal two-way nondenaturing polyacrylamide gel (details in Materials and Methods) and staining with SYBR Gold to visualize RNA, we observed a single band for protein-free RNA that migrated toward the anode (there was no visible fluorescence for the lane that only contained protein). For samples including constant [protein] and varied [RNA], we observed a new band migrating at a slower rate (also toward the anode) that increased in intensity with each increase in RNA concentration (as well as an increase in intensity of the band representing free RNA, see Figure 2).

**Figure 2.**
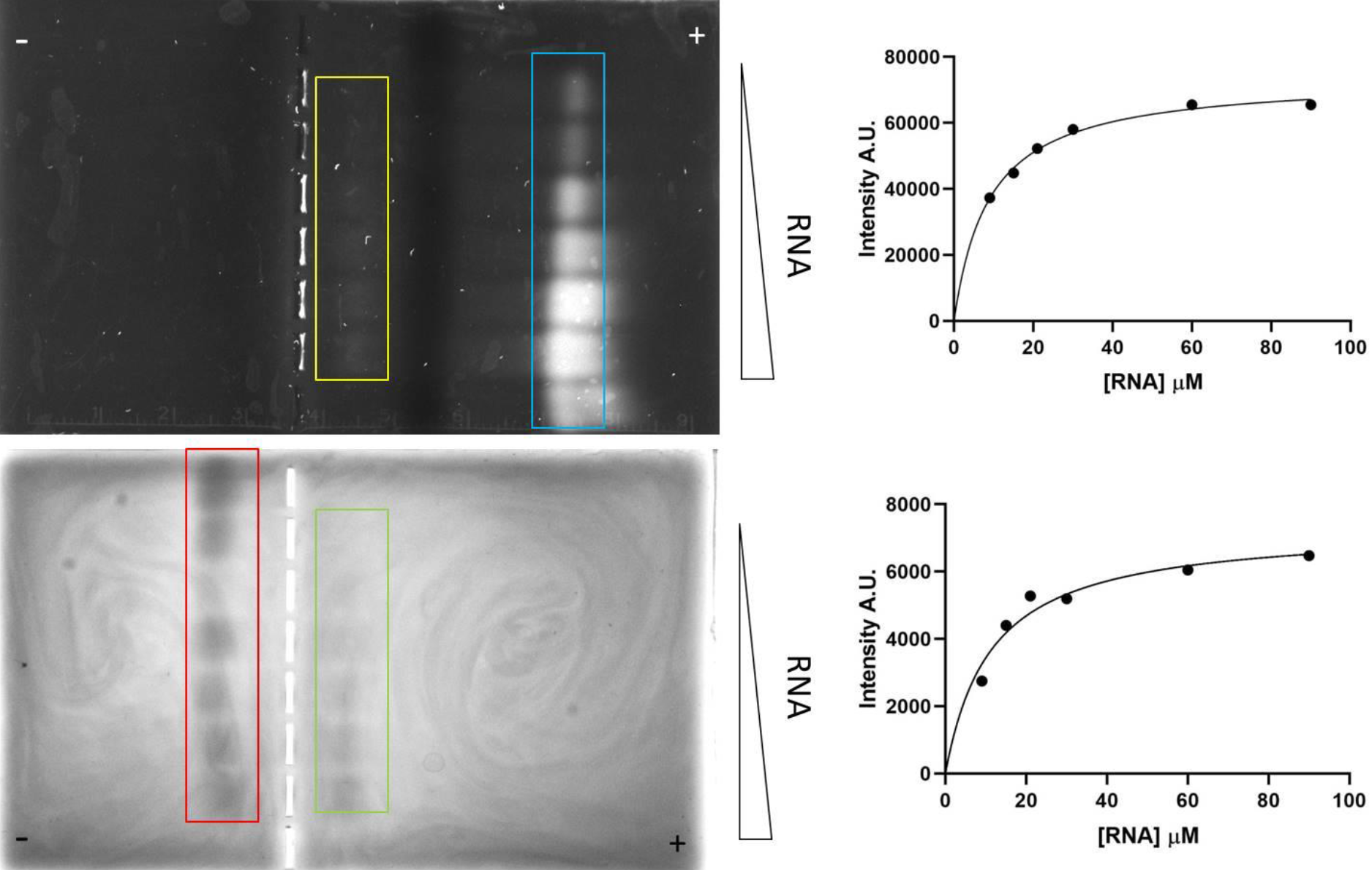
Non-denaturing Horizontal PAGE in which splicing factor p14 (30 μM) was titrated with a 13-nucleotide RNA representing the intron’s branch region (9 – 90 μM). **Top**: after staining with SyBr Gold. Cyan box: free RNA. Yellow box: protein-RNA complex. **Bottom**: After staining with Coomassie Blue. Red box: Free protein. Green box: protein-RNA complex. Ratios of RNA:protein range from 0.3:1 to 3:1 to 30μM amount of protein. Fitting of the best-fit curve indicated values of K_d_ = 8.9 ± 0.8 μM (SYBR Gold) and 6.6 ± 1.9 μM (Coomassie Blue). Additional experimental details in Materials and Methods.

After staining the same gel with Coomassie Blue, we observed that the band for the RNA-free protein p14 (pI 9.4) migrated toward the cathode. Each sample with increasing amounts of RNA, from 9 to 90 μM, was associated with an increase in the intensity of the shifted protein band in the identical location of the shifted RNA band visualized by SYBR gold, as well as a fractional decrease in the band of the unbound protein. We then quantified the intensities of each band for each of the two staining procedures with UVP VisionWorks software (uvp.com/visionworks) and plotted the data of intensity vs. μM RNA (plot shown next to the stained gel from which data were obtained). For the SYBR Gold data, the data were best fit with a line describing a 1:1 stoichiometry and a Kd = 8.9 ± 1.9 μM (Figure 2, top); for the Coomassie Blue data for the same gel, the best fit curve indicated 1:1 stoichiometry and a Kd = 6.6 ± 1.3 μM (also on Figure 2). These data indicate that, independent of the staining method used, the interaction affinity between RNA and protein are very similar.

We also performed the assays in the reverse “direction”, *i.e*. titrating p14 protein (9 – 90 μM) into constant RNA (30 μM), with ratios of protein:RNA ranging from 0.3:1 to 3:1 (Figure 3). Increased intensity of the protein-RNA complex moving toward the anode was observed upon addition of increased amounts of protein and a single band for the RNA-free protein or the protein free-RNA moving toward the anode and towards the cathode respectively. The equilibrium dissociation constants obtained after nonlinear fitting (6.3 ± 0.9 μM (SYBR Gold) and 4.9 ± 1.6 μM (Coomassie Blue) were very similar to the ones observed before. Our results indicate very similar values for each experimental configuration.

**Figure 3.**
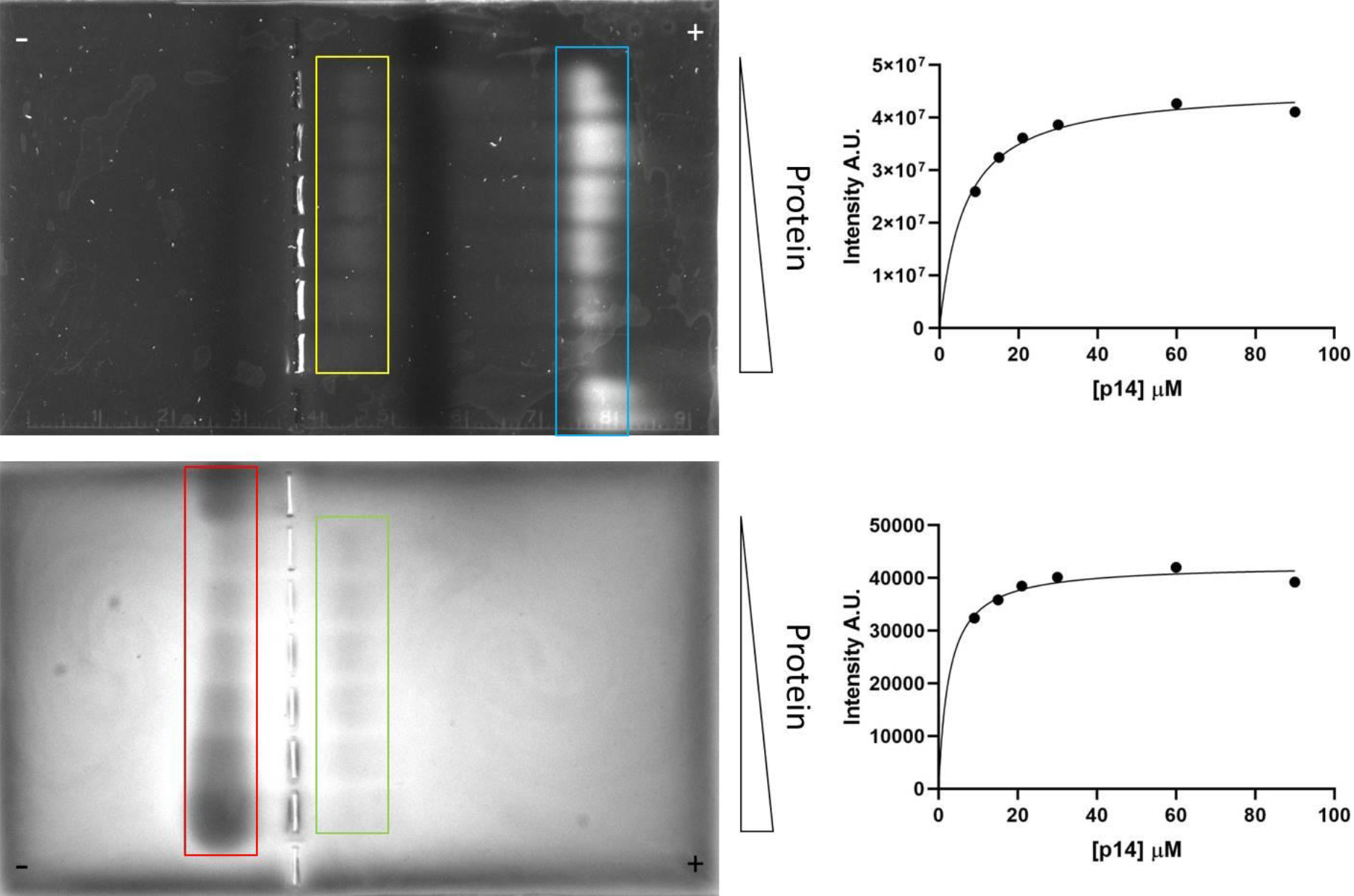
Non-denaturing Horizontal PAGE in which the 13-nucleotide RNA (30 μM) was titrated with protein splicing factor p14 (9 – 90 μM). **Top**: after staining with SyBr Gold. Cyan box: free RNA. Yellow box: protein-RNA complex. **Bottom**: After staining with Coomassie Blue. Red box: free protein. Green box: protein-RNA complex. Ratios of protein:RNA range from 0.3:1 to 3:1 to 30 μM amount of RNA. Fitting of the best-fit curve indicated values of K_d_ = 6.3 ± 0.9 μM (SYBR Gold) and 4.9 ± 1.6 μM (Coomassie Blue). Additional experimental details in Materials and Methods.

## Discussion

Binding and specificity in the interaction of RBPs with their RNA targets are crucial in the biophysical characterization of biomolecular complexes, such as those associated with the spliceosome or the ribosome, where multiple proteins are important in the assembly and function of the particles. A systematic study of such interactions would benefit from a straightforward method to assay affinity and specificity, such as that provided by EMSA measurements. A problem with these methods is the need to pre-label the RNA target and the difficulty to detect and quantify the free proteins and protein-RNA complexes for RBPs with high pI which migrate toward the cathode during electrophoresis. In an attempt to address this issue we develop a simple strategy to monitor protein-RNA complexes migration in a simple horizontal PAGE. There are many approaches to measuring biomolecular affinity, each with advantages and disadvantages/limitations: EMSA with labels; optical/spectroscopic methods (NMR, CD, fluorescence polarization/anisotropy); surface plasmon resonance (Biacore); calorimetric methods (ITC and Microscale thermophoresis) provide information about thermodynamic parameters. The method describe here allowed us to determine that presence of the affinity His-tag commonly used to ease the purification of recombinant proteins affects the binding properties of the protein and it was necessary to remove it for the desired assays (data not shown).

## Conclusions

Protein-RNA interactions can be characterized by a diverse number of methods including EMSA, a ubiquitous and powerful technique easily adapted to a variety of systems [5]. The method described here allows to detect and to quantitate the K_d_ for the interaction of an RNA binding protein regardless of its pI. The method allows screening binding conditions for the biophysical characterization of protein/RNA interactions. Although the sensitivity or our method is far less than other methods, the simplicity and time-saving ability of the method allows characterizing protein-RNA systems when enough amounts of biomolecules are available. Conditions like gel percent, binding buffers, continuous buffer pH, voltage and length of running for a particular protein-RNA system can be easily optimized.

## Acknowledgments

Authors thank past and present graduate and undergraduate students in our lab for testing and optimizing experimental conditions for several RNA-protein interactions using the method described in this communication.

## References

1. Hellman, L. M., and Fried, M. G. (2007) Electrophoretic mobility shift assay (EMSA) for detecting protein-nucleic acid interactions. Nat. Protoc. 2, 1849–1861.

2. Ryder, S. P., Recht, M. I., and Williamson, J. R. (2008) Quantitative analysis of protein-RNA interactions by gel mobility shift. Methods Mol. Biol. 488, 99–115.

3. Castello, A., Fischer, B., Eichelbaum, K., Horos, R., Beckmann, B. M., Strein, C., Davey, N. E., Humphreys, D. T., Preiss, T., Steinmetz, L. M., Krijgsveld, J., and Hentze, M. W. (2012) Insights into RNA Biology from an Atlas of Mammalian mRNA-Binding Proteins. Cell 149, 1393–1406.

4. Perea, W., Schroeder, K.T., Bryant, A.N., and Greenbaum, N.L. (2016). Interaction between the Spliceosomal Pre-mRNA Branch Site and U2 snRNP Protein p14. Biochemistry 55, 629–632.

5. Dong, W. H., Wang, T. Y., Wang, F., and Zhang, J. H. (2011) Simple, Time-Saving Dye Staining of Proteins for Sodium Dodecyl Sulfate–Polyacrylamide Gel Electrophoresis Using Coomassie Blue. PLoS One 6(8), e22394.

6. Ramanathan M., Porter, D. F., and Khava, P. A. (2019) Methods to Study RNA-Protein Interactions. Nat. Methods 16, 225–234.

